# TreeGazer: Prospecting Protein Sequence-Function Landscapes via Phylogenetic Structure

**DOI:** 10.64898/2026.05.14.725301

**Authors:** Sebastian Porras, Samuel Davis, Oscar Paredes Trujillo, Patrick Diep, Gerhard Schenk, Mikael Bodén

## Abstract

Building diverse and informative protein sequence datasets is critical for understanding how function varies across sequence space. Because only a small fraction of sequences in a dataset can typically be experimentally characterised, strategies for selecting what sequences to characterise should maximise the information gained from each experiment. Here, we present TreeGazer, a phylogeny-informed framework that combines Bayesian optimisation with the topology of a tree to guide sequence selection. TreeGazer balances exploitation of sequences predicted to exhibit favourable properties against exploration of regions higher model uncertainty. Unlike existing approaches that apply Bayesian optimisation for sequence selection, TreeGazer does not rely on black-box models and instead uses latent representations of property distributions that are directly tied to phylogenetic structure. Modelling properties in this way enables biologically interpretable predictions and uncertainty estimates. Across two simulated selection campaigns, TreeGazer consistently selected sequences that produced datasets more representative of the underlying property distribution than alternative strategies that used protein language models. TreeGazer also performed effectively in low-data settings, where tree-guided selection enabled accurate identification of functional transitions across clades. TreeGazer can be run on conventional laptop computers while still providing equivalent or superior performance to embedding-based approaches. These results demonstrate that phylogenetic structure is a powerful and underutilised prior for guiding informative sequence selection.

## 1 Introduction

Understanding the relationship between a protein’s sequence and its function is inherently complex. The vast combinatorial nature of sequence space means that many mutations are neutral and have very little affect, obscuring the mutations that meaningfully alter function. Characterising this landscape experimentally is costly, and so any campaign aimed at understanding the sequence-function relationship depends critically on selecting a maximally informative set of sequences [Arnold, 2019, Zhu et al., 2022]. This challenge is compounded in low-data regimes, where coverage of sequence space is sparse and the risk of building a biased, fragmented picture of the landscape is high.

Bayesian optimisation [Shahriari et al., 2016] is a principled approach to this problem, explicitly balancing *exploitation* of regions near sequences with desirable properties against *exploration* of regions with high model uncertainty. A model’s prediction can be uncertain when annotations are sparse or where data are conflicting. However, the utility of Bayesian optimisation depends heavily on the underlying sequence representation. Approaches based on protein language model (PLM) embeddings [Yang et al., 2025] or kernel functions [Romero et al., 2013] have been previously applied, but both carry a shared limitation: the representations are high-dimensional and difficult to interpret, making it hard to track functional changes across these representation spaces [Notin et al., 2023].

High-throughput strategies such as deep mutational scanning (DMS) [Fowler et al., 2010] can partially circumvent the sparsity of experimental annotations by densely sampling a small region of sequence space, but this comes at the cost of breadth; mutations are typically limited to a handful of residues, and the resulting datasets do not capture the functional diversity that exists across orthologous proteins [Copp et al., 2018, Krco et al., 2023]. For example, because of epistasis, the effect of mutating two or more interacting positions in a sequence is different to what would be expected from combining the individual effect of each mutation [Starr et al., 2017]. Without diverse datasets, epistasis can be difficult to detect and can obscure underlying biological mechanisms explaining function [Anderson et al., 2015, Kondrashov and Kondrashov, 2015, Brookes et al., 2022]. Although emerging technologies aim to overcome these limitations [Muir et al., 2024, Long et al., 2025, Diep et al., 2026], the need for interpretable methods to guide sequence selection remains.

Phylogenetic trees offer a compelling alternative foundation for sequence selection. Rather than mapping sequences into an opaque embedding space, a tree provides an explicitly interpretable structure in which evolutionary distance serves as a proxy for *functional* divergence. By grouping sequences into clades — monophyletic groups sharing a common ancestor — a phylogeny amplifies evolutionary signal while reducing the influence of noisy, inconsequential sequence variation [Thomas et al., 2003, Ortiz-Velez et al., 2024]. Conservation within clades and distinction between them can reveal the sequence determinants of functional diversification, providing an interpretable framework for understanding the sequence-function relationship.

Ancestral sequence reconstruction (ASR) expands on the phylogenetic framework by predicting the sequences of internal nodes on a tree. The characterisation of ancestors provides information intermediate to extant sequences and hence confers an enhanced ability to identify key determinants of functional variation across a tree [Thornton, 2001]. Ancestors at branch points corresponding to major evolutionary events are of particular interest, as they likely represent functional junctures in the protein’s history.

Despite the richness of the information encoded in a phylogeny, sequence selection from trees remains largely guided by biochemical intuition rather than formal methods. Estimating how likely a functional property is to be retained over a given evolutionary distance is non-trivial, and no principled framework exists for leveraging tree structure to systematically explore sequence space in a data-driven way. Evolutionary biochemists often must rely on expert knowledge to make predictions about the properties of a given protein sequence by observing the branch lengths in combination with available experimental data for related proteins which may be distantly positioned on a tree. This process of selecting sequence is therefore often fraught with bias and the process is difficult to describe and reproduce. We ask: given a small number of experimental observations, can a method that explicitly couples the evolutionary structure of a phylogenetic tree to a latent representation of protein properties guide the construction of datasets that faithfully approximate the true distribution of those properties across sequence space? We were particularly motivated by low-data regimes, where the cost of biased or redundant sequence selection is highest, and where the interpretability and biological grounding of a tree-based approach offer the greatest advantage over existing methods. Neglecting exploration in this regime leads to a biased and fragmented view of the sequence-function landscape which is the very outcome that principled sequence selection is designed to avoid.

## 2 Methods

### 2.1 Modelling protein properties with TreeGazer

TreeGazer was developed to harness information encoded by a phylogeny to estimate values of arbitrary properties at uncharacterised nodes and quantify the associated statistical uncertainty. TreeGazer identifies information-deficient regions of a tree under a Bayesian optimisation framework which balances exploitation and exploration. Unlike many existing approaches that aim to optimise specific properties or rely on black-box models [Wu et al., 2019, Hie et al., 2020, Yang et al., 2025], TreeGazer models properties with clear assumptions and requires no advanced computational expertise.

TreeGazer builds on the GRASP-Suite [Foley et al., 2022], which represents phylogenetic trees as Bayesian networks (BNs). Unlike most property prediction methods that use a protein’s sequence, TreeGazer requires only a phylogenetic tree and known annotation values corresponding to at least one node. Given a fixed, rooted, and bifurcating phylogenetic tree, TreeGazer constructs a BN mirroring its topology to model discrete latent variables, the state space of which is assumed to capture unobserved functional modes evolving along branches (Figure 1). The latent state distribution of a child node is therefore conditional only on the state of its parent and is a function of branch length, analogous to sequence character reconstruction in ASR. In a biochemical context, transitions between latent evolutionary modes represent significant mutations, driven by selection for the value of interest.

**Figure 1:**
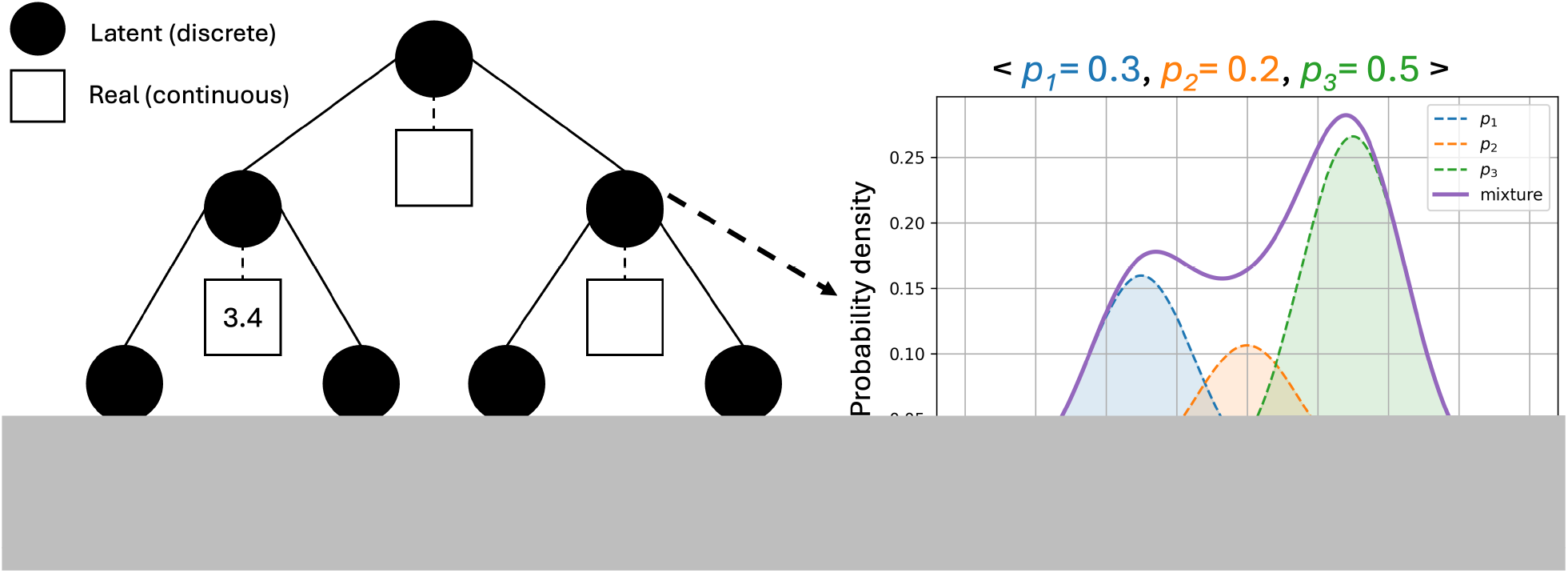
Depiction of a Phylo-Bayesian network. Discrete latent nodes (circles) mimic the structure of the phylogenetic tree, while continuous, real nodes (squares) are fixed with known values for a given property (for example, a kinetic parameter). Users can specify any number of latent states, with each latent state mapping to a Gaussian distribution for which parameters are learned from the data via EM. Predictions of properties and the corresponding uncertainty of this prediction can be obtained by creating a Gaussian mixture from each latent Gaussian, weighted by the marginal probability of the latent state.

Each element of the state space – or evolutionary *mode* – corresponds to a Gaussian density over values of the modelled property, and together these modes define a mixture model over the property’s distribution across the tree. Continuous-valued nodes are conditioned on each latent node and correspond to probability densities of the property’s value for the associated extant or ancestral sequence. Gaussian features allow for normally distributed variation between nodes for which the latent state mode is identical, capturing stochastic variation between homologous proteins where background mutations contribute indirectly to small changes in the trait value. The latent state space and associated Gaussian feature parameters are assumed to be constant across the tree, such that their underlying encoding of the property of interest is inherent for the given protein family. Once the size of the latent sample space, *k*, is chosen, the set of associated Gaussian parameters,

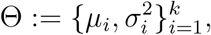

are learned by expectation maximisation [Dempster et al., 1977], using known experimental values as input.

A generalised form of the Jukes-Cantor model for nucleotide substitution [Jukes and Cantor, 1969] is used to model transitions between latent states. Conditional probabilities between nodes are functions of time specified by the branch length, in which the time to change latent states is exponentially distributed with a rate, *α*. As with the original Jukes-Cantor model that assumes that all nucleotide substitutions are equally likely, the likelihood of transitioning to any other latent state is equal. When generalised to *k* latent states, the time dependent substitution model is specified by the following transition probability matrix:

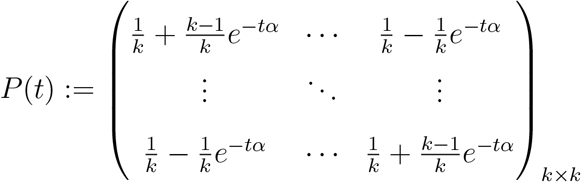

where diagonal values represent the probability of remaining in the same state, and off-diagonal values represent equally likely transitions to another latent state.

Maximum likelihood inference of latent states for unannotated nodes are performed via variable elimination (VE) as described by Dechter [1998]. VE sums over all possible assignments to each latent node. The resulting marginal probability distribution, {*p*_1_, …, *p*_*k*_}, at a latent node specifies a continuous probability density at the real node of interest. The inference of the node’s continuous feature value as a mixture of Gaussian features weighted by the respective latent state probabilities is defined as

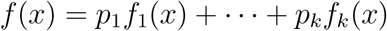

where *p*_*i*_ represents the probability of the latent state assigned to the element *i* and *f*_*i*_ is the corresponding Gaussian density. The standard deviation of this mixture provides a natural measure of prediction confidence.

TreeGazer generates interpretable visualisations compatible with Interactive Tree Of Life (ITOL) [Letunic and Bork, 2024], allowing data to be directly mapped onto the phylogenetic tree (Figure 2) for easy identification of statistical uncertainty. Learning and inference can be performed in under one minute on standard desktop hardware for the adenylate kinase (ADK) [Muir et al., 2024] example in Figure 2 (175 leaves) which negates the requirement for high performance hardware.

**Figure 2:**
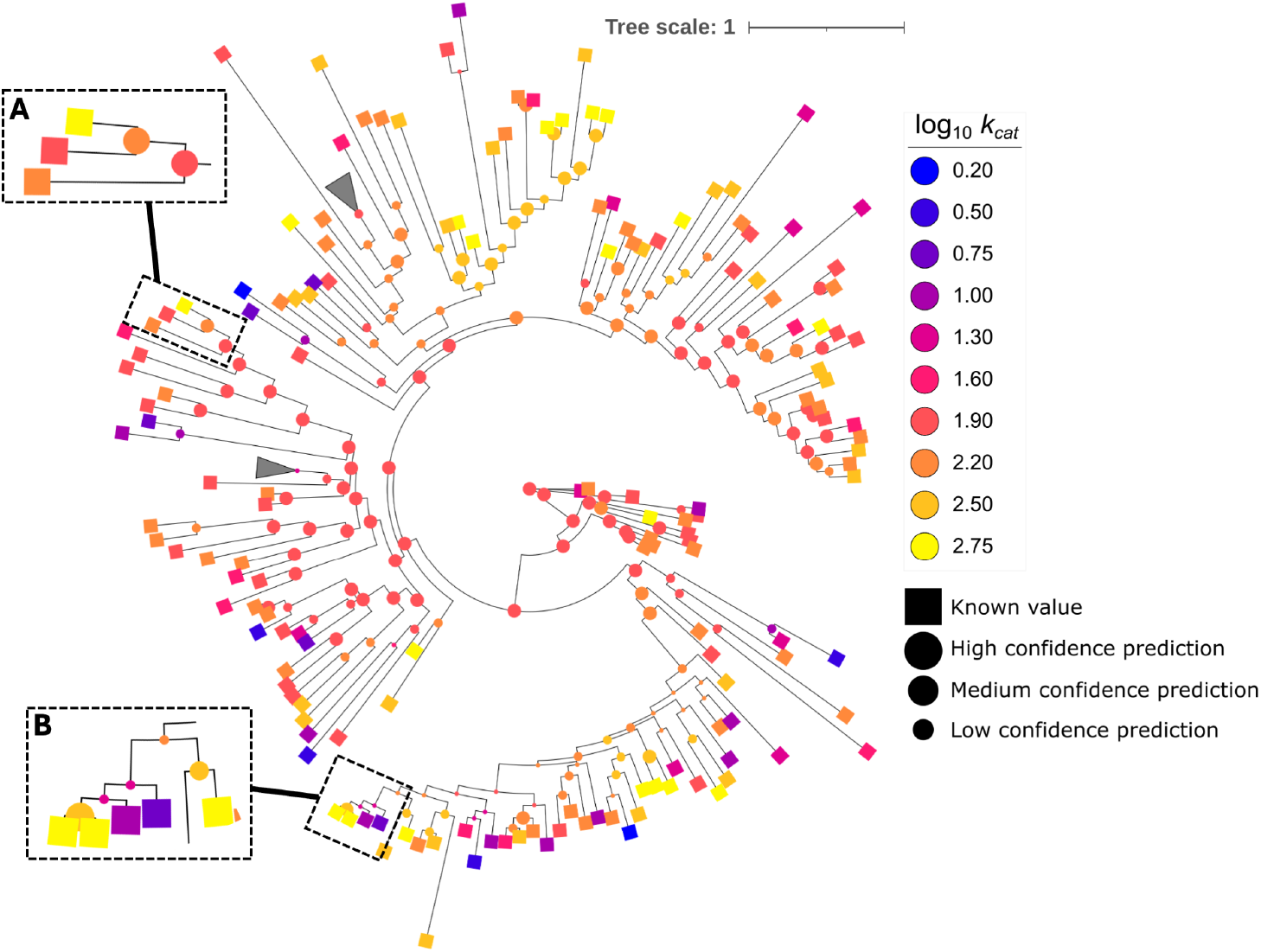
Example Interactive Tree of Life visualisation produced by TreeGazer. Squares represent experimentally-derived values and circles represent predicted features. Node colour corresponds to value, and circle size represents model confidence, with larger circles indicating higher confidence. **A:** The set of extants have relatively similar property values (2.2-2.75 log_10_ *k*_*cat*_) with short branches resulting in high confidence predictions. **B:** Extant property values span from 0.2 to 2.75 log_10_ *k*_*cat*_ but also have short branches which makes it easy to say that the prediction is uncertain.

It is worth noting that TreeGazer is highly dependent on the quality of the input tree topology; careful consideration should be taken when constructing the tree to ensure it is representative of the underlying sequence space. The multiple sequence alignment used to construct the tree should balance the breadth of taxonomic coverage with alignment quality. Including as many sequences as possible while avoiding introducing too much ambiguity is essential, as the alignment has a substantial affect on the resulting tree [Talavera and Castresana, 2007, Ross et al., 2022]. Input sequences should also be carefully curated for technical errors, as the expansion of sequence databases has increased the prevalence of transcription errors and miscalled genomic events such as frameshifts, insertions, and deletions [Thomson et al., 2022]. Finally, trees should be inferred using maximum likelihood or Bayesian methods rather than distance-based approaches, as probabilistic approaches have been shown to produce more accurate topologies [Rannala and Yang, 1996].

### 2.2 Balancing information gain against maximisation of favourable properties

Effectively optimising a property or understanding how that property evolved requires diverse and informative samples rather than greedily sampling near the best performing sequence. TreeGazer addresses this need by applying a Bayesian optimisation framework [Shahriari et al., 2016] to guide sequence selection in a statistically informed manner. The selection of candidates strikes a balance between *exploration* (characterising regions of high model uncertainty to maximise information gain) and *exploitation* (selecting the most likely favourable proteins) by considering the uncertainty associated with model predictions. TreeGazer implements the widely used upper confidence bound (UCB) acquisition function. For a node *x* with mean *µ* and standard deviation *σ* determined from the Gaussian mixture, the UCB acquisition function is defined as:

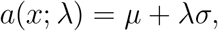

with *λ* influencing the degree of exploration. Sequence selection then proceeds by identifying the node *x* that maximises the acquisition function *a*.

### 2.3 Datasets used for evaluation

To evaluate the effectiveness of TreeGazer at selecting informative samples we used two datasets with only extant property annotations and one dataset that contains ancestral annotations to highlight the flexibility of the TreeGazer framework. For the extant datasets we drew on two publicly available datasets from the ADK [Muir et al., 2024] and lanmodulin [Diep et al., 2026] protein families. These datasets are unique as they contain a set of orthologous sequences that are comprehensively annotated with properties determined by cutting edge, high-throughput assays, providing the breadth of functional and sequence diversity necessary to construct a meaningful phylogenetic tree.

The ADK dataset from Muir et al. [2024] contains 175 ADK orthologs with catalytic parameters that provide comprehensive coverage of *K*_*M*_ and *k*_*cat*_ values. The lanmodulin dataset provided by Diep et al. [2026] consists of 616 extant sequences which were annotated with metal selectivity for the entire lanthanide set including Y^III^ and La^III^ to Lu^III^, excluding Pm^III^ and Sc^III^. For each of the ADK and lanmodulin datasets a tree and a corresponding substitution model was inferred using IQ-TREE2 (version 2.1.2) [Minh et al., 2020] with 1000 Ultrafast bootstrap replicates and ModelFinder [Kalyaanamoorthy et al., 2017], respectively.

For the ancestral dataset, we previously performed ASR on the class I ketol-acid reductoisomerases (KARIs) protein family with the aim of improving the catalytic efficiency of these enzymes for the production of (R)-2,3-dihydroxy-3-isovalerate (DHIV) in the presence of isobutanol [Trujillo et al., 2025]. That study identified an ancestral variant that displayed increased activity in the presence of isobutanol. Here we sought to determine whether TreeGazer could identify other sequences which could reasonably expected to display similar isobutanol tolerance. The tree used in that study contained 716 extant KARIs with catalytic parameters and isobutanol tolerance assays for 8 ancestors and 2 extants. In addition to the previously reported annotations, we performed isobutanol tolerance assays for three further ancestors and two extants in this work.

The isobutanol tolerance assays were performed using an Agilent Cary 60 UV-Vis spectrophotometer. The activity of every KARI was measured by monitoring the oxidation of NADH at 340 nm, a reaction that is coupled to the conversion of (S)-2-acetolactate to DHIV. The isobutanol tolerance assays were performed using a reaction buffer at pH 8 in a total volume of 1 mL. The reaction solution was composed of 0.2 mM NAD(P)H, 1 µM of enzyme, 10 mM MgCl_2_, 100 mM HEPES, 5 mM (S)-2-acetolactate (saturating concentration) and increasing concentrations of isobutanol (0 - 10% v/v). The assays were performed at 25 °C without pre-incubation. The influence of isobutanol was evaluated by comparing the residual activity of the KARIs in the presence of isobutanol (2 – 10 % v/v) with the activity at 0% v/v isobutanol which we refer to as relative activity.

### 2.4 Simulation of a selection campaign

The comprehensive ADK and lanmodulin extant datasets were used to simulate a selection campaign, evaluating how effectively TreeGazer selects candidates that reveal information about the underlying property distribution. The campaign proceeded as follows:

1. A random 20% of the data was designated as the training dataset; the remaining 80% of data comprised the test dataset, from which candidate sequences were drawn at each iteration. This procedure was repeated 30 times with independent random initialisations.
2. At each iteration, kernel density estimation (KDE) [Pedregosa et al., 2011] was used to approximate two property distributions: one over the current training dataset and one over the combined training and test datasets, the latter serving as an estimate of the true property distribution. The Jensen-Shannon distance [Lin, 1991] — a symmetric, bounded measure of dissimilarity between probability distributions — was then computed between these two distributions. A more informative selection campaign is expected to reduce this distance more rapidly as selected candidates are added to the training data.
3. Based on the training data, each model ranked the candidate sequences in the test dataset according to its acquisition function (see Section 2.5).
4. The highest-ranking candidate was transferred from the test dataset to the training data, and steps 2-4 were repeated until the test dataset was exhausted.

### 2.5 Models compared against TreeGazer

To contextualise the performance of TreeGazer, we compared it against three baseline models spanning greedy [Edmonds, 1971] and Bayesian optimisation strategies. The two greedy baselines select candidates solely by predicted property value, without accounting for model uncertainty, while the Bayesian approaches balance exploitation with exploration via the UCB acquisition function.

As a phylogeny-informed greedy baseline, we implemented weighted distance estimate (WDE), which predicts the property value of an unknown candidate *X* as a weighted mean of inverse evolutionary distances to annotated nodes. Formally:

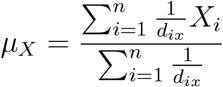

where *µ*_*X*_ is the predicted property value, *d*_*ix*_ is the evolutionary distance between a known sequence *i* and the candidate sequence *X*, and *X*_*i*_ is the property value at a known sequence *i*. The sequence with the greatest predicted value, *µ*_*X*_, was selected at each iteration.

The second greedy baseline approach used mean-pooled ProtT5-XL-UniRef50 [Elnaggar et al., 2022] embeddings to represent sequences, with property values predicted by k-nearest neighbour (k-NN) scoring, specifically the average of the three nearest neighbours by cosine distance in the embedding space. As with WDE, the candidate with the greatest predicted value was selected at each iteration.

For the Bayesian optimisation baseline, ProtT5-XL-UniRef50 embeddings were used to train a Radial basis function kernel Gaussian process (RBF-GP) using GPyTorch [Gardner et al., 2018]. Candidate sequences were ranked using the UCB acquisition function (*λ* = 5) with the mean and standard deviation drawn from the posterior predictive distribution.

TreeGazer was initialised with three latent variable states with tied variance across the Gaussian densities. Three latent states were chosen because using greater than three states produced multiple Gaussian densities with the same mean, indicating that additional states were non-informative. Gaussian density parameters were fit to the training data via EM until convergence, and property distributions for test sequences were obtained by marginal inference over the corresponding tree nodes. Candidates were ranked using UCB (*λ* = 5), with the mean and standard deviation taken from the resulting Gaussian mixture.

## 3 Results and Discussion

### 3.1 TreeGazer consistently selects informative sequences

When performing the simulated selection campaign, we compared TreeGazer against three other approaches for a total of 30 replicates for each dataset. For the ADK dataset, TreeGazer consistently gained more information about the true distribution than other methods for both *k*_*cat*_/*K*_*M*_ (Figure 3A) and *k*_*cat*_ (Figure 3B). Early in each selection campaign, all methods performed comparably. However, once the training dataset exceeded 100 samples, the mean Jensen-Shannon distance for TreeGazer was substantially lower than for the other methods.

**Figure 3:**
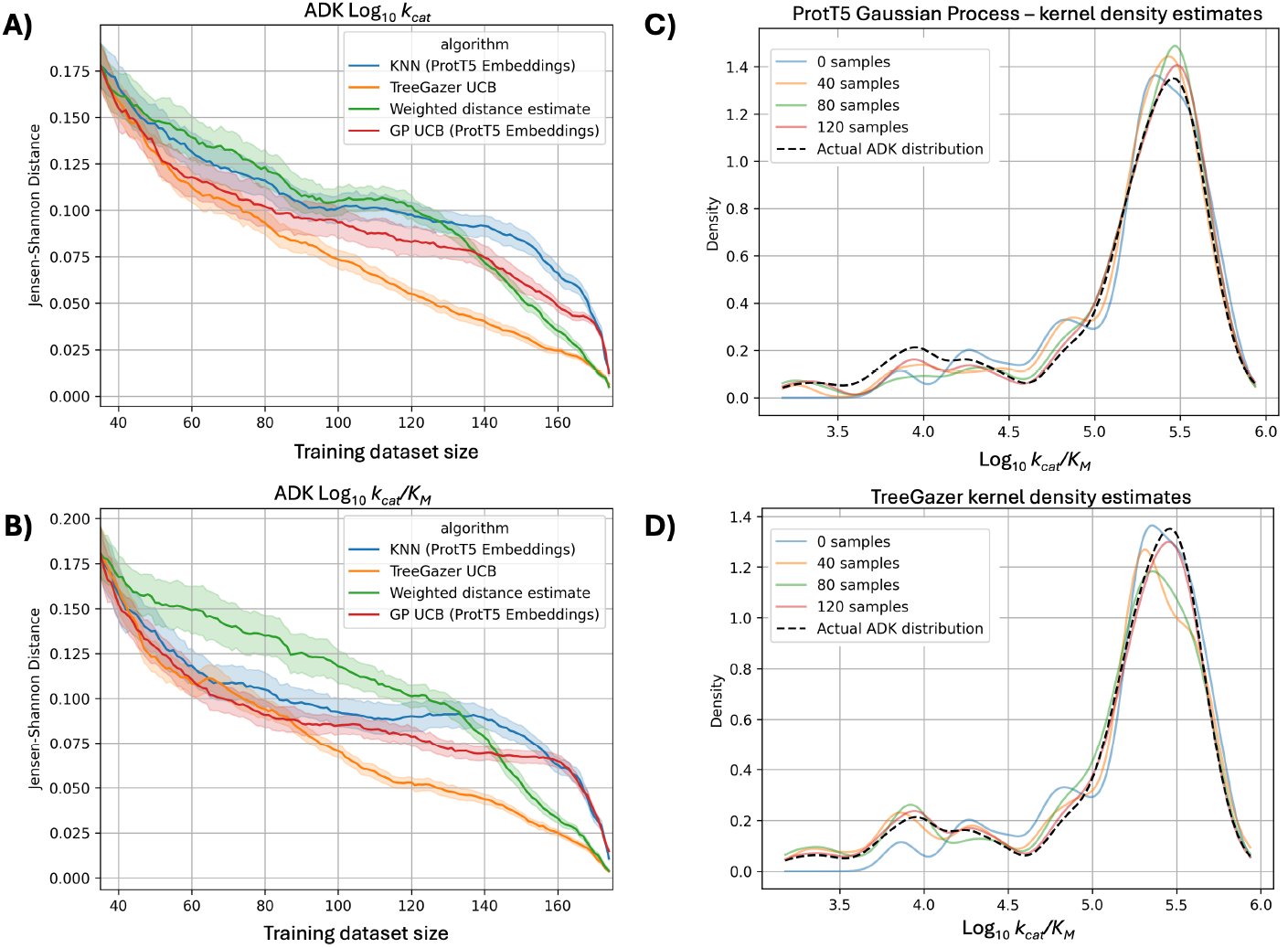
**A & B**: Results of selection campaigns on log_10_ *k*_*cat*_ (A) and log_10_ *k*_*cat*_/*K*_*M*_ (B). With 30 random trials, each model was provided with 20% of the data and tasked with recommending one additional sequence. This sequence was then added to the training data and this process was repeated until all data points were revealed. At each iteration, the Jensen-Shannon distance between the true distribution and the training data distribution was measured by approximating each distribution with kernel density estimation. For both properties, TreeGazer consistently gains information and provides a more accurate estimate of the true distribution as measured by the Jensen-Shannon distance. **C & D**: Distributions approximated by kernel density estimates at different training dataset sizes for the ProtT5 Gaussian process (C) and TreeGazer (D) for log_10_ *k*_*cat*_/*K*_*M*_. Panel C demonstrates how the ProtT5 Gaussian process assigns much more probability to the distribution mean and undersamples values below 4.5 log_10_ *k*_*cat*_/*K*_*M*_ compared to TreeGazer shown in panel D which explains why TreeGazer had relatively lower Jensen-Shannon distances.

Both the k-NN and RBF-GP models plateaued in information gain when the training dataset contained between 85 and 140 samples, whereas TreeGazer made consistent gains throughout. Visualisations of the KDE distributions (Figure 3C and 3D) demonstrate how TreeGazer sampled low density (but information-rich) sections of the property distribution more effectively than other models. TreeGazer approximated the true distribution more closely in regions where annotations for *k*_*cat*_/*K*_*M*_ values between 3.5 and 5 log M^*−*1^ s^*−*1^ were sparse (Figure 3D). In contrast, the RBF-GP model favoured samples near the distribution mean (Figure 3C) and modelled the tail regions poorly.

The underperformance of the embedding-based approaches in the selection campaign is likely attributable to the nature of the representations themselves. Although embeddings contain some phylogenetic information [Tule et al., 2025], they also encode a multitude of other biochemical and structural information, making it difficult to assess why certain embeddings are considered similar. This conflation of biological information in embeddings may discourage sampling variants that are informative for understanding functional divergence and specialisation for a particular property of interest. This drawback is particularly apparent when the goal is exploration rather than optimisation.

While the ADK dataset consists of a set of sequences evenly dispersed across evolutionary sequence space, the lanmodulin dataset contains clusters of highly similar sequences separated by very long branches as highlighted by the phylogenetic distance matrix in Figure 4 and the tree itself in Figure S1. This topology resulted in the selection campaign proceeding very differently from the ADK case. Rather than consistently gaining information, all methods unevenly sampled the tree space and became trapped in local optima, repeatedly drawing from clades of highly similar sequences (Figure 4A and Figure 4B). As a result, approximated distributions were skewed towards the property values of these clades, increasing the Jensen-Shannon distance from the true distribution. TreeGazer also suffered from biased sampling, but the severity was substantially less than other methods (Figure 4C). The long branches separating clades increased model uncertainty at distance nodes, providing TreeGazer with a mechanism for escaping optima unlike the embedding-based approaches that lack an explicit representation of evolutionary distance.

**Figure 4:**
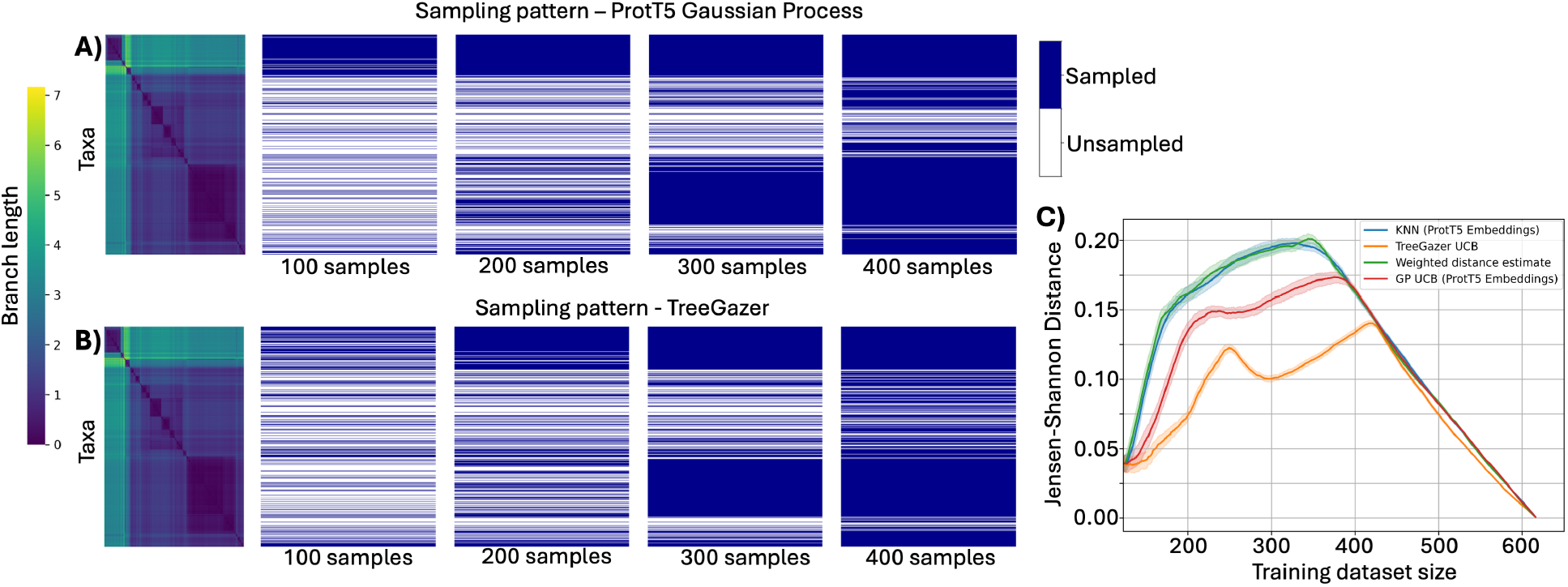
**A & B**: Extants sampled by the ProtT5 Gaussian process (A) and TreeGazer (B) at successive stages of the selection campaign. The rows are clustered according to phylogenetic distance, with a blue row indicating the extant has been included in the training data. **C**: Results of the selection campaign across all replicates for the lanmodulin protein family. With 30 random trials, each model was provided with 20% of the data and tasked with recommending one additional sequence. This sequence was then added to the training data and this process was repeated until all data points were revealed. At each iteration, the Jensen-Shannon distance between the true distribution and the training data distribution was measured by approximating each distribution with kernel density estimation.

### 3.2 TreeGazer provides quantitative evidence for biological intuition

TreeGazer can help direct wet-lab resources towards characterising sequences (ancestral or extant) that provide functional insights with as few experiments as possible. We recently applied ASR to the KARI family to identify potential candidates for a cell-free enzyme cascade for the conversion of glucose to isobutanol [Trujillo et al., 2025], identifying a unique ancestral KARIs variant that showed a positive correlation between the concentration of isobutanol activity. We sought to pinpoint the ancestral node at which isobutanol tolerance was gained, with the aim of investigating the biochemical mechanisms underpinning this tolerance and determining whether other sequences within the clade would display similar properties.

Because TreeGazer makes predictions and estimates uncertainty according to tree structure, branch lengths and the grouping of annotations implied by topology become important sources of information when interpreting model outputs. We trained TreeGazer with three latent states on the KARI dataset (two extants and eight ancestors) to model activity at 8% isobutanol and obtain a statistical estimate of where the gain of isobutanol activation occurred. We hypothesised that this gain isobutanol activation occurred along the branch between N28 and N79. This hypothesis was supported by TreeGazer’s prediction that N28 would not exhibit isobutanol activation, whereas N78 should retain elevated activity (Figure 5A). TreeGazer also predicted a general decay in the level of isobutanol activation from the highly isobutanol tolerant N79 towards the leaves.

**Figure 5:**
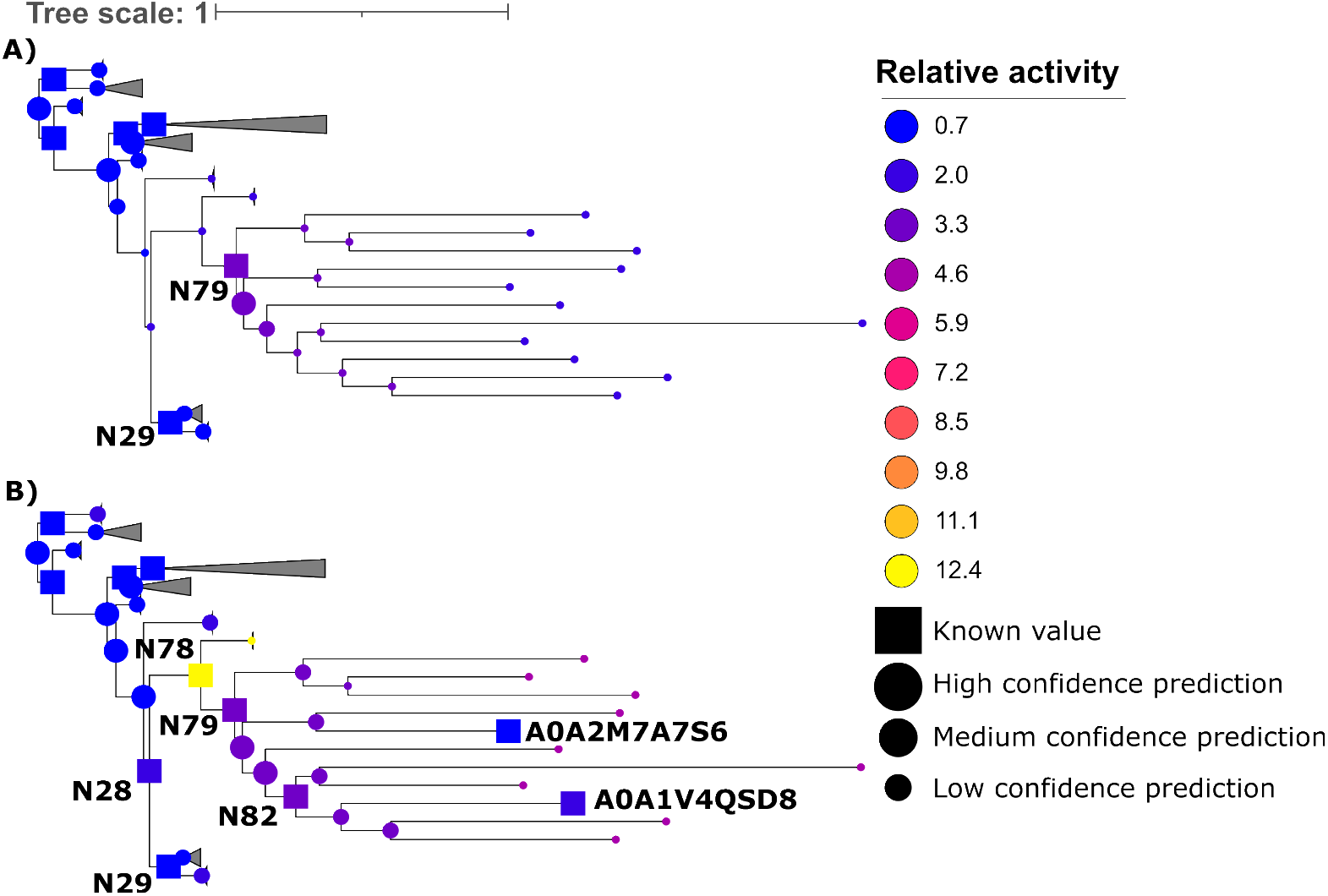
**A**: Initial TreeGazer predictions for relative activity at 8% isobutanol prior to further characterisation of sequences within the clade. Relative activity is the ratio of enzyme activity in the presence of 8% isobutanol to activity with 0% isobutanol. TreeGazer predicted higher relative activity in the presence of isobutanol for all members of the clade below N79 and the direct parent of this ancestor, N78. **B**: TreeGazer predictions for relative activity at 8% isobutanol following the additional characterisation of three ancestors and two extants. N82 and the two extants demonstrated moderate increased activity in the presence of isobutanol and N78 was more than 12 times more active.

To verify these predictions and test whether isobutanol activation extended to other members of the clade, we characterised ancestors N28, N78 and N82 along with two extants (UniProt IDs A0A2M7A7S6 and A0A1V4QSD8). TreeGazer correctly identified the boundary of isobutanol activation. N28 showed decreased relative activity (0.86) to the control with no isobutanol present, while N82 and the two extants displayed moderate activation, but not to the same extent as N79. Ancestor N78 was an outlier, exhibiting activity more than 12-fold greater than its own no-isobutanol baseline (Figure 5B). Having confirmed the distribution of isobutanol tolerance across this clade and identified the point of transition, these findings provide a foundation for investigating the molecular mechanisms contributing to this phenotype.

Beyond guiding sequence selection, TreeGazer predictions also offer a means of assessing whether a tree’s topology adequately captures how a property is distributed across sequence space and by extension, whether the property is well-suited to tree-based modelling at all. Where evolutionary relationships closely track functional variation, as in proteins are under consistent selective pressure, the tree topology will correlate well with property variation. For example, a protein common to a group of thermophilic bacteria may consistently be under selective pressure for high thermostability. The variation in thermostability between the protein variants would reflect the environmental niches or amino acid preferences of the source organism and we expect TreeGazer would perform reliably. In contrast, for promiscuous protein families capable of acting on a multitude of substrates [Babbitt et al., 1996], shifts in selection pressure may be poorly reflected by the tree topology and TreeGazer’s predictions will be correspondingly uncertain. Rather than being a limitation, this uncertainty is itself informative. Annotations that cluster closely on the tree but exhibit unexpectedly high statistical uncertainty signal either that the tree topology inadequately represents the sequence space, or that the property is not effectively represented using the topology of a tree. In either case, this diagnostic capacity makes TreeGazer useful for developing a more rigorous understanding of how a given property can best be modelled. Even for researchers familiar with evolutionary biochemistry, approaches like TreeGazer can provide phylogenetic intuition for both experienced practitioners and newcomers to the field.

## 4 Conclusion

Understanding how protein function varies across a family of orthologous sequences remains a fundamental challenge in molecular biology. Because experimental characterisation is costly, sequences selected for assay should maximise what is learned about the true distribution of functional properties across sequence space. By simulating a selection campaign, we demonstrated that tree-informed approaches consistently selected sequences that built more accurate approximations of a property distribution per iteration that PLM-based strategies across two comprehensively annotated phylogenetic datasets with distinct characteristics. In practical terms, a wet-lab scientist using TreeGazer would arrive at a representative picture of how a property varies across a protein family with fewer experiments than with existing approaches.

This advantage stems from TreeGazer’s explicit use of tree structure as a prior for functional variation. Rather than relying on embeddings that may obscure evolutionary signal, TreeGazer ties inference directly to evolutionary relationships and quantifies uncertainty in a biologically interpretable way. In the KARI case study, TreeGazer produced credible predictions of how isobutanol tolerance distributes across a clade even in an extremely low-data setting. TreeGazer correctly identified the point of functional transition and guided experimental validation that confirmed and extended these findings. TreeGazer’s uncertainty estimates also serve as a diagnostic tool for assessing whether a given property is well-suited to tree-based modelling and can provide insights into the evolutionary basis of functional variation.

TreeGazer offers equivalent or superior performance to embedding-based approaches at a fraction of the computational cost with far greater interpretability. As comprehensively annotated orthologous datasets become more widely available, phylogeny-informed methods will become an increasingly important complement to sequence-based approaches for navigating the sequence-function landscape.

## 6 Author contributions statement

S.P, S.D and M.B developed the software, designed and performed research and wrote the paper. P.D contributed with data and P.D, O.P.T and G.S contributed to writing the paper. O.P.T performed wet-lab experiments.

## References

D. W. Anderson, A. N. McKeown, and J. W. Thornton. Intermolecular epistasis shaped the function and evolution of an ancient transcription factor and its DNA binding sites. eLife, 4:e07864, June 2015. ISSN 2050-084X. doi: 10.7554/eLife.07864. URL https://doi.org/10.7554/eLife.07864.

F. H. Arnold. Innovation by Evolution: Bringing New Chemistry to Life (Nobel Lecture). Angewandte Chemie International Edition, 58(41):14420–14426, Oct. 2019. ISSN 1433-7851. doi: 10.1002/anie.201907729.

P. C. Babbitt, M. S. Hasson, J. E. Wedekind, D. R. J. Palmer, W. C. Barrett, G. H. Reed, I. Rayment, D. Ringe, G. L. Kenyon, and J. A. Gerlt. The Enolase Superfamily: A General Strategy for Enzyme-Catalyzed Abstraction of the α-Protons of Carboxylic Acids. Biochemistry, 35(51):16489–16501, Jan. 1996. ISSN 0006-2960. dio: 10.1021/bi9616413. URL https://doi.org/10.1021/bi9616413.

D. H. Brookes, A. Aghazadeh, and J. Listgarten. On the sparsity of fitness functions and implications for learning. Proceedings of the National Academy of Sciences, 119(1): e2109649118, Jan. 2022. doi: 10.1073/pnas.2109649118. URL https://www.pnas.org/doi/full/10.1073/pnas.2109649118.

J. N. Copp, E. Akiva, P. C. Babbitt, and N. Tokuriki. Revealing Unexplored Sequence-Function Space Using Sequence Similarity Networks. Biochemistry, 57(31):4651–4662, Aug. 2018. ISSN 0006-2960. dio: 10.1021/acs.biochem.8b00473.

R. Dechter. Bucket Elimination: A Unifying Framework for Probabilistic Inference. In M. I. Jordan, editor, Learning in Graphical Models, pages 75–104. Springer Netherlands, Dordrecht, 1998. ISBN 978-94-011-5014-9. doi: 10.1007/978-94-011-5014-9_4.

P. Dempster, N. M. Laird, and D. B. Rubin. Maximum Likelihood from Incomplete Data Via the EM Algorithm. Journal of the Royal Statistical Society: Series B (Methodological), 39(1):1–22, Sept. 1977. ISSN 0035-9246. dio: 10.1111/j.2517-6161.1977.tb01600.x.

P. Diep, C. S. Madsen, W. Choi, Z. Dong, C. S. Kang-Yun, P. F. V. Uychoco, J. A. Seidel, S. A. Eaton, Y. Jiao, J. A. Cotruvo, and D. M. Park. A family portrait of lanmodulin selectivity for enhanced rare-earth separations. Nature Chemical Biology, 22(5):829–839, May 2026. ISSN 1552-4469. dio: 10.1038/s41589-026-02176-3.

J. Edmonds. Matroids and the greedy algorithm. Mathematical Programming, 1(1):127–136, Dec. 1971. ISSN 1436-4646. dio: 10.1007/BF01584082.

Elnaggar, M. Heinzinger, C. Dallago, G. Rehawi, Y. Wang, L. Jones, T. Gibbs, T. Feher, C. Angerer, M. Steinegger, D. Bhowmik, and B. Rost. ProtTrans: Toward Understanding the Language of Life Through Self-Supervised Learning. IEEE Transactions on Pattern Analysis and Machine Intelligence, 44(10):7112–7127, Oct. 2022. ISSN 1939-3539. dio: 10.1109/TPAMI.2021.3095381.

G. Foley, A. Mora, C. M. Ross, S. Bottoms, L. Sützl, M. L. Lamprecht, J. Zaugg, A. Essebier, B. Balderson, R. Newell, R. E. S. Thomson, B. Kobe, R. T. Barnard, L. Guddat, G. Schenk, J. Carsten, Y. Gumulya, B. Rost, D. Haltrich, V. Sieber, E. M. J. Gillam, and M. Bodén. Engineering indel and substitution variants of diverse and ancient enzymes using Graphical Representation of Ancestral Sequence Predictions (GRASP). PLOS Computational Biology, 18(10):e1010633, Oct. 2022. ISSN 1553-7358. dio: 10.1371/journal.pcbi.1010633.

D. M. Fowler, C. L. Araya, S. J. Fleishman, E. H. Kellogg, J. J. Stephany, D. Baker, and S. Fields. High-resolution mapping of protein sequence-function relationships. Nature Methods, 7(9):741–746, Sept. 2010. ISSN 1548-7105. dio: 10.1038/nmeth.1492.

J. Gardner, G. Pleiss, K. Q. Weinberger, D. Bindel, and A. G. Wilson. GPyTorch: Blackbox matrix-matrix gaussian process inference with GPU acceleration. In S. Bengio, H. Wallach, H. Larochelle, K. Grauman, N. Cesa-Bianchi, and R. Garnett, editors, Advances in neural information processing systems, volume 31. Curran Associates, Inc., 2018.

Hie, B. D. Bryson, and B. Berger. Leveraging Uncertainty in Machine Learning Accelerates Biological Discovery and Design. Cell Systems, 11(5):461–477.e9, Nov. 2020. ISSN 2405-4712, 2405-4720. dio: 10.1016/j.cels.2020.09.007. URL https://www.cell.com/cell-systems/abstract/S2405-4712(20)30364-1.

T. H. Jukes and C. R. Cantor. Evolution of protein molecules. In Mammalian Protein Metabolism, pages 21–132. Academic Press, Jan. 1969. ISBN 978-1-4832-3211-9. doi: 10.1016/B978-1-4832-3211-9.50009-7.

S. Kalyaanamoorthy, B. Q. Minh, T. K. F. Wong, A. von Haeseler, and L. S. Jermiin. ModelFinder: fast model selection for accurate phylogenetic estimates. Nature Methods, 14(6):587–589, June 2017. ISSN 1548-7105. dio: 10.1038/nmeth.4285.

A. Kondrashov and F. A. Kondrashov. Topological features of rugged fitness landscapes in sequence space. Trends in Genetics, 31(1):24–33, Jan. 2015. ISSN 0168-9525. dio: 10.1016/j.tig.2014.09.009. URL https://www.sciencedirect.com/science/article/pii/S0168952514001577.

S. Krco, S. J. Davis, P. Joshi, L. A. Wilson, M. Monteiro Pedroso, A. Douw, C. J. Schofield, P. Hugenholtz, G. Schenk, and M. T. Morris. Structure, function, and evolution of metallo-β-lactamases from the B3 subgroup—emerging targets to combat antibiotic resistance. Frontiers in Chemistry, Volume 11-2023, 2023. ISSN 2296-2646. dio: 10.3389/fchem.2023.1196073.

Letunic and P. Bork. Interactive Tree of Life (iTOL) v6: recent updates to the phylogenetic tree display and annotation tool. Nucleic Acids Research, 52(W1):W78–W82, July 2024. ISSN 0305-1048. dio: 10.1093/nar/gkae268.

J. Lin. Divergence measures based on the Shannon entropy. IEEE Transactions on Information Theory, 37(1):145–151, Jan. 1991. ISSN 1557-9654. dio: 10.1109/18.61115.

Y. Long, A. Mora, F.-Z. Li, E. Gürsoy, K. E. Johnston, and F. H. Arnold. LevSeq: Rapid Generation of Sequence-Function Data for Directed Evolution and Machine Learning. ACS Synthetic Biology, 14(1):230–238, Jan. 2025. doi: 10.1021/acssynbio.4c00625.

Q. Minh, H. A. Schmidt, O. Chernomor, D. Schrempf, M. D. Woodhams, A. von Haeseler, and R. Lanfear. IQ-TREE 2: New Models and Efficient Methods for Phylogenetic Inference in the Genomic Era. Molecular Biology and Evolution, 37(5):1530–1534, May 2020. ISSN 0737-4038. dio: 10.1093/molbev/msaa015.

F. Muir, G. P. R. Asper, P. Notin, J. A. Posner, D. S. Marks, M. J. Keiser, and M. M. Pinney. Evolutionary-Scale Enzymology Enables Biochemical Constant Prediction Across a Multi-Peaked Catalytic Landscape. bioRxiv, page 2024.10.23.619915, Jan. 2024. doi: 10.1101/2024.10.23.619915.

P. Notin, A. Kollasch, D. Ritter, L. van Niekerk, S. Paul, H. Spinner, N. Rollins, A. Shaw, R. Orenbuch, R. Weitzman, J. Frazer, M. Dias, D. Franceschi, Y. Gal, and D. Marks. ProteinGym: Large-Scale Benchmarks for Protein Fitness Prediction and Design. In Oh, T. Naumann, A. Globerson, K. Saenko, M. Hardt, and S. Levine, editors, Advances in Neural Information Processing Systems, volume 36, pages 64331–64379. Curran Associates, Inc., 2023.

N. Ortiz-Velez, J. Sukumaran, R. Rouzbehani, and S. T. Kelley. AutoPhy: Automated phylogenetic identification of novel protein subfamilies. PLOS ONE, 19(1):e0291801, Jan. 2024. doi: 10.1371/journal.pone.0291801.

F. Pedregosa, G. Varoquaux, A. Gramfort, V. Michel, B. Thirion, O. Grisel, M. Blondel, P. Prettenhofer, R. Weiss, V. Dubourg, J. Vanderplas, A. Passos, D. Cournapeau, M. Brucher, M. Perrot, and E. Duchesnay. Scikit-learn: Machine learning in python. Journal of Machine Learning Research, 12(85):2825–2830, 2011. URL http://jmlr.org/papers/v12/pedregosa11a.html.

Rannala and Z. Yang. Probability distribution of molecular evolutionary trees: A new method of phylogenetic inference. Journal of Molecular Evolution, 43(3):304–311, Sept. 1996. ISSN 1432-1432. dio: 10.1007/BF02338839. URL https://doi.org/10.1007/BF02338839.

P. A. Romero, A. Krause, and F. H. Arnold. Navigating the protein fitness landscape with Gaussian processes. Proceedings of the National Academy of Sciences, 110(3):E193–E201, Jan. 2013. doi: 10.1073/pnas.1215251110.

M. Ross, G. Foley, M. Boden, and E. M. J. Gillam. Using the Evolutionary History of Proteins to Engineer Insertion-Deletion Mutants from Robust, Ancestral Templates Using Graphical Representation of Ancestral Sequence Predictions (GRASP). In F. Magnani, C. Marabelli, and F. Paradisi, editors, Enzyme Engineering: Methods and Protocols, pages 85–110. Springer US, New York, NY, 2022. ISBN 978-1-0716-1826-4. doi: 10.1007/978-1-0716-1826-4_6. URL https://doi.org/10.1007/978-1-0716-1826-4_6.

Shahriari, K. Swersky, Z. Wang, R. P. Adams, and N. De Freitas. Taking the Human Out of the Loop: A Review of Bayesian Optimization. Proceedings of the IEEE, 104(1): 148–175, Jan. 2016. ISSN 1558-2256. dio: 10.1109/JPROC.2015.2494218.

T. N. Starr, L. K. Picton, and J. W. Thornton. Alternative evolutionary histories in the sequence space of an ancient protein. Nature, 549(7672):409–413, Sept. 2017. ISSN 1476-4687. dio: 10.1038/nature23902. URL https://www.nature.com/articles/nature23902. Number: 7672.

G. Talavera and J. Castresana. Improvement of Phylogenies after Removing Divergent and Ambiguously Aligned Blocks from Protein Sequence Alignments. Systematic Biology, 56 (4):564–577, Aug. 2007. ISSN 1063-5157. dio: 10.1080/10635150701472164. URL https://doi.org/10.1080/10635150701472164.

P. D. Thomas, M. J. Campbell, A. Kejariwal, H. Mi, B. Karlak, R. Daverman, K. Diemer Muruganujan, and A. Narechania. PANTHER: A Library of Protein Families and Subfamilies Indexed by Function. Genome Research, 13(9):2129–2141, Sept. 2003. ISSN 1088-9051, 1549-5469. dio: 10.1101/gr.772403. Company: Cold Spring Harbor Laboratory Press Distributor: Cold Spring Harbor Laboratory Press Institution: Cold Spring Harbor Laboratory Press Label: Cold Spring Harbor Laboratory Press.

R. E. S. Thomson, S. E. Carrera-Pacheco, and E. M. J. Gillam. Engineering functional thermostable proteins using ancestral sequence reconstruction. Journal of Biological Chemistry, 298(10), Oct. 2022. ISSN 0021-9258, 1083-351X. dio: 10.1016/j.jbc.2022.102435. URL https://www.jbc.org/article/S0021-9258(22)00878-X/abstract.

J. W. Thornton. Evolution of vertebrate steroid receptors from an ancestral estrogen receptor by ligand exploitation and serial genome expansions. Proceedings of the National Academy of Sciences, 98(10):5671–5676, May 2001. doi: 10.1073/pnas.091553298.

O. P. Trujillo, G. Foley, S. Porras, G. Joyce, A. Douw, S. Tule, H. Ehlert, L. Asser, U. Adhikary, S. Davis, D. Hine, M. Marzo, K. Evans, J. Ley, V. Sieber, L. Guddat, M. Boden, and G. Schenk. Engineering stable and efficient ketol-acid reductoisomerases for industrial biotransformations using ancestral sequence reconstruction., Dec. 2025. ISSN: 2693-5015.

S. Tule, G. Foley, and M. Bodén. Do protein language models learn phylogeny? Briefings in Bioinformatics, 26(1):bbaf047, Jan. 2025. ISSN 1477-4054. dio: 10.1093/bib/bbaf047.

Z. Wu, S. B. J. Kan, R. D. Lewis, B. J. Wittmann, and F. H. Arnold. Machine learning-assisted directed protein evolution with combinatorial libraries. Proceedings of the National Academy of Sciences, 116(18):8852–8858, Apr. 2019. doi: 10.1073/pnas.1901979116. URL https://www.pnas.org/doi/full/10.1073/pnas.1901979116.

J. Yang, R. G. Lal, J. C. Bowden, R. Astudillo, M. A. Hameedi, S. Kaur, M. Hill, Y. Yue, and F. H. Arnold. Active learning-assisted directed evolution. Nature Communications, 16(1):714, Jan. 2025. ISSN 2041-1723. dio: 10.1038/s41467-025-55987-8.

B. Zhu, D. Wang, and N. Wei. Enzyme discovery and engineering for sustainable plastic recycling. Trends in Biotechnology, 40(1):22–37, Jan. 2022. ISSN 0167-7799. dio: 10.1016/j.tibtech.2021.02.008.

