## Supplemental information for "TreeGazer: Prospecting Protein Sequence-Function Landscapes via Phylogenetic Structure"

### 1 Supplementary

TreeGazer source code and JAR file are available at https://github.com/bodenlab/bnkit. Analysis scripts and data are available at https://github.com/SebPorras/TreeGazer_paper_resources.

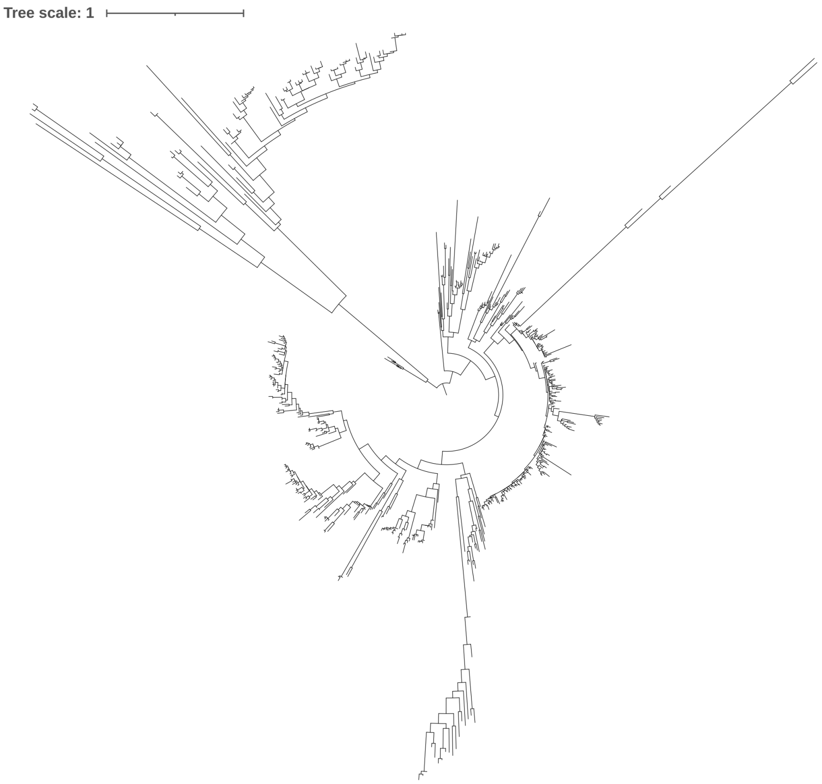


Figure S1: The phylogenetic tree of lanmodulin sequences. The tree contained 616 extant sequences. The sequences were aligned using MAFFT before tree inference was performed with IQ-TREE2.
